# The abundance and diversity of West Nile virus mosquito vectors in two Regional Units of Greece during the onset of the 2018 transmission season

**DOI:** 10.1101/735522

**Authors:** Marina Bisia, Claire L Jeffries, Ioanna Lytra, Antonios Michaelakis, Thomas Walker

**Author notes:** authors contributed equally to the work (Marina Bisia, Claire L Jeffries).

## Abstract

**Background:** West Nile virus (WNV) is a zoonotic arbovirus of great medical and veterinary importance, threatening the health of humans and equines worldwide. Mosquitoes belonging to the *Culex* (*Cx*.) *pipiens* complex are major vectors but numerous other mosquito species have also been implicated as vectors of WNV. Due to variations in blood-feeding behaviour, the different biotypes and hybrids of *Cx. pipiens* influence the transmission of WNV, from enzootic cycles (between mosquitoes and birds), to spill-over transmission to humans and equines.

**Methods:** In this study, mosquitoes were collected and analysed from two regional units (RUs) of Greece with reported cases of WNV within the past 4 years; Palaio Flairo and Argolida (in Attica and Peloponnese regions, respectively). Collections using different types of mosquito surveillance traps were undertaken in May-June 2018 during the early period of the WNV transmission season.

**Results:** A total of 1062 mosquitoes were collected, with Biogents Sentinel traps (BG traps) collecting both a greater number of mosquitoes across all species and *Cx. pipiens* complex individuals than Centres for Disease Control miniature light traps (CDC traps) or Heavy Duty Encephalitis Vector Survey traps (EVS traps). Identification of collected mosquitoes (using both morphological keys and molecular barcoding) confirmed the presence of additional species including *Aedes (Ae.) albopictus, Ae. caspius* and *Culiseta (Cs.) longiareolata*. The prevalence of *Cx. pipiens* biotypes in the RU of Palaio Faliro was 54.5% *pipiens* type, 20.0% *molestus* type and 25.5% hybrids. In the RU of Argolida, the collection comprised 68.1% *pipiens* type, 8.3% *molestus* type and 23.6% hybrids. Screening individual unfed female mosquitoes for WNV (molecular xenomonitoring) resulted in detection in three females of the *pipiens* type and in one hybrid; all collected from the RU of Argolida.

**Conclusions:** As hybrids play an important role in spill-over transmission of WNV to humans and equines, these findings highlight the importance of undertaking entomological surveillance programs incorporating molecular xenomonitoring at the onset of the transmission season to provide an early warning system for health authorities aiming to prevent WNV outbreaks in Greece.

## Background

West Nile virus (WNV) is an arbovirus belonging to the Japanese encephalitis group within the *Flavivirus* genus (*Flaviviridae* family) and is the most widespread virus belonging to this genus [1–4]. Natural transmission of WNV mainly occurs in enzootic cycles between birds and competent ornithophilic mosquito vectors, with avian species being the principal maintenance and amplifying hosts of WNV as many species develop sufficient viremia for onward transmission. This allows transmission to continue where competent mosquitoes are present in a specific area under suitable environmental conditions [5]. Additionally, spill-over transmission can occur when competent vectors feed on humans or horses. During natural transmission these mammalian species are considered dead-end hosts since they cannot sustain sufficient viraemia for further vector-borne transmission. However, infection in humans does pose a transmission risk due to the possibility of iatrogenic transmission through blood and tissue donations, in addition to the possibility of intrauterine transmission or WNV being passed on through breast milk [4]. Blood and tissue donor screening is essential in areas where WNV is endemic [6,7] and currently no human vaccination is available, however, vaccination of horses has been shown to reduce clinical disease in this species [8,9].

WNV was first isolated in 1937 from a woman with febrile illness in the West Nile district of Uganda [10]. WNV has caused numerous recent outbreaks in North America and Europe leading to major concern for human and animal health [3,11]. In North America, the majority of arboviral encephalitis cases are attributable to WNV [12]. Although ~80% of human WNV infections are asymptomatic, a broad clinical spectrum can result ranging from a mild flu-like illness in ~20% of infected individuals (West Nile fever) to severe neurological disease through infection of the central nervous system (<1% of infected individuals) that can lead to death from meningitis, encephalitis, and acute flaccid paralysis [13,14]. The high proportion of asymptomatic infections highlights that the number of human cases demonstrating overt disease, or discovered through laboratory testing, are just the ‘tip of the iceberg’ of the actual number of viral infections occurring within a population. Furthermore, these spill-over infections in humans are likely to be far less frequent compared to the amount of enzootic transmission occurring between mosquitoes and avian species. This emphasises the high value of surveillance in the monitoring and prevention of major outbreaks.

The introduction and spread of WNV in Europe is thought to have been driven by migratory birds [15–18]. WNV resulted in sporadic human cases from the mid-1990s [19] with the first large outbreak occurring in Romania with 393 hospitalised cases and 17 deaths [19,20]. From 2010, the European Center for Disease Control (ECDC) have monitored WNV cases in the European Union and neighbouring countries and publishes weekly epidemiological reports [21]. In Greece, WNV was first detected in the summer of 2010 in the central Macedonia Region near the city of Thessaloniki, in the northern part of the country [22,23]. This outbreak included 262 probable and confirmed cases of WNV infection of which 197 were neuroinvasive cases and 35 deaths [24]. In 2011 WNV was found in both humans and horses; detected from clinical and laboratory surveillance techniques [25]. In the following years, cases of WNV in humans and animals were reported in central Greece and in the Attica Region but there were no reported cases in 2015 or 2016. In 2017, WNV re-emerged in southern Greece and in 2018 there were 311 laboratory confirmed human cases, resulting in 47 deaths, showing a marked increase over 2017, with only 48 confirmed cases and 5 deaths [21,25]. Historical data of human cases with neurological disease in Greece from 2010 until present show that cases increase in August (the peak month in the transmission season) and the largest case numbers were reported in August 2010 [24–26].

There have been over 60 species of mosquitoes in the USA implicated as potential WNV vector species [4]. Seven of these species occur in Europe and have been tested for WNV susceptibility including members of the *Culex (Cx.) pipiens* complex, *Ae. albopictus* and *Ae. (Ocherlotatus) caspius* [27]. *Cx. pipiens* has two behaviourally different biotypes, *pipiens* and *molestus*, which can form hybrids and their feeding behaviours influence their role in local transmission of WNV. The *pipiens* biotype is an important species for the enzootic WNV transmission cycle given its preference to feed on birds [28]. The *molestus* biotype and hybrids are implicated in the spill-over transmission of WNV from avian hosts to humans due to the opportunistic feeding behaviour of the *molestus* biotype [28,29]. Temperature has been shown to experimentally increase WNV transmission rates of the *pipiens* and hybrid biotypes but have no effect on the *molestus* biotype [30]. In order to better understand the complexity of WNV transmission, entomological surveys for arboviral surveillance can be undertaken to determine both the presence of potential mosquito vectors and provide evidence for WNV circulation through virus detection in field-caught mosquitoes (molecular xenomonitoring). Here we report the results of an entomological survey undertaken in two Regional Units (RUs) of Greece (Palio Faliro in Attica region and Argolida in Peloponnese region) where WNV outbreaks have previously been recorded. We determined the prevalence of the *Cx. pipiens* biotypes (*pipiens*, *molestus* and hybrids) in each sampling location and female mosquitoes were screened for the presence of WNV to determine whether there was any evidence of virus circulation in the two RUs.

## Methods

### Mosquito collections

The study was carried out in 2 RUs within the Attica and Peloponnese regions of Greece, with three sampling locations selected from within each RU, and three trapping sites within each sampling location (**Fig. 1, Table 1**). Locations for trapping in the RU of Palaio Faliro were classified as urban, whereas those in the RU of Argolida were rural. In each sampling location, three different traps (trapping sites) were operating for 24 hours, three times per week. Trapping occurred over a six-week period (May-June 2018) during the start of the WNV transmission season (based on previous historical data obtained from ECDC [31]). A 3×3 Latin square design [32] was applied at each site to minimize confounding factors. Traps were placed more than 100m from each other and rotated every 24 hours between selected positions. Three different trap types were used in each site; Biogent sentinel (BG) traps, Heavy duty Encephalitis Vector Survey (EVS) traps and Centers for Disease Control miniature light (CDC) traps. Dry ice was used as an attractant in all traps with approximately 2 kg/trap per 24 hours. Mosquitoes were collected every 24 hours, killed on dry ice and stored at −80°C. Morphological keys were used to identify individuals to species or species complex level [33] and female mosquitoes were classified as unfed (no evidence of blood in their abdomen), blood fed or gravid. Individual mosquitoes were then placed in RNAlater (Invitrogen) to preserve RNA for downstream molecular analysis.

**Fig. 1.**
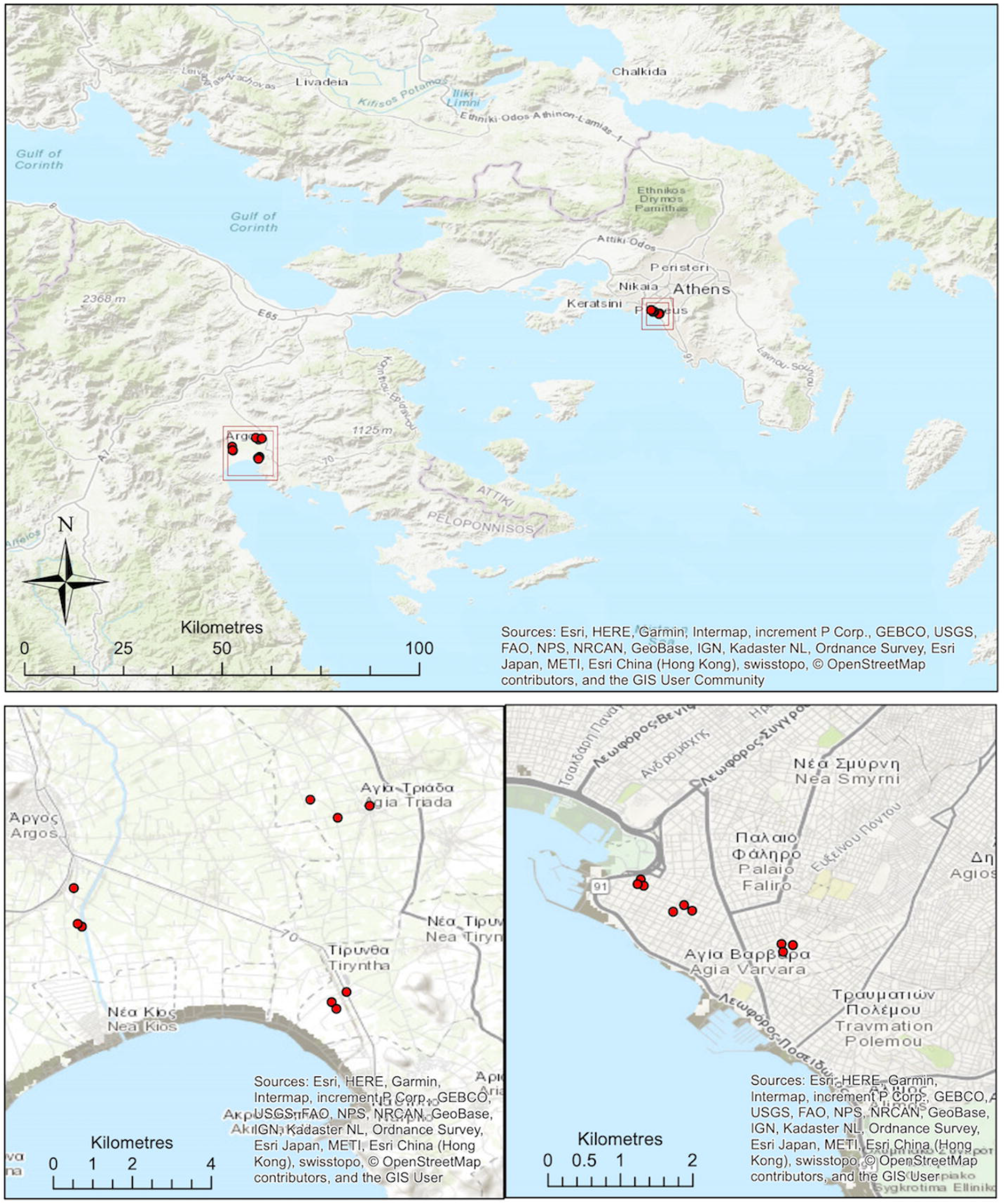
Locations of collection sites within Regional Unit of Palaio Faliro in Attica Region (lower right) and Regional Unit of Argolida in Peloponnese region (lower left). Maps constructed in ArcMap 10.5 (Esri, ArcGIS), using World Topographic Basemap and GPS coordinates from trap locations.

**Table 1.**
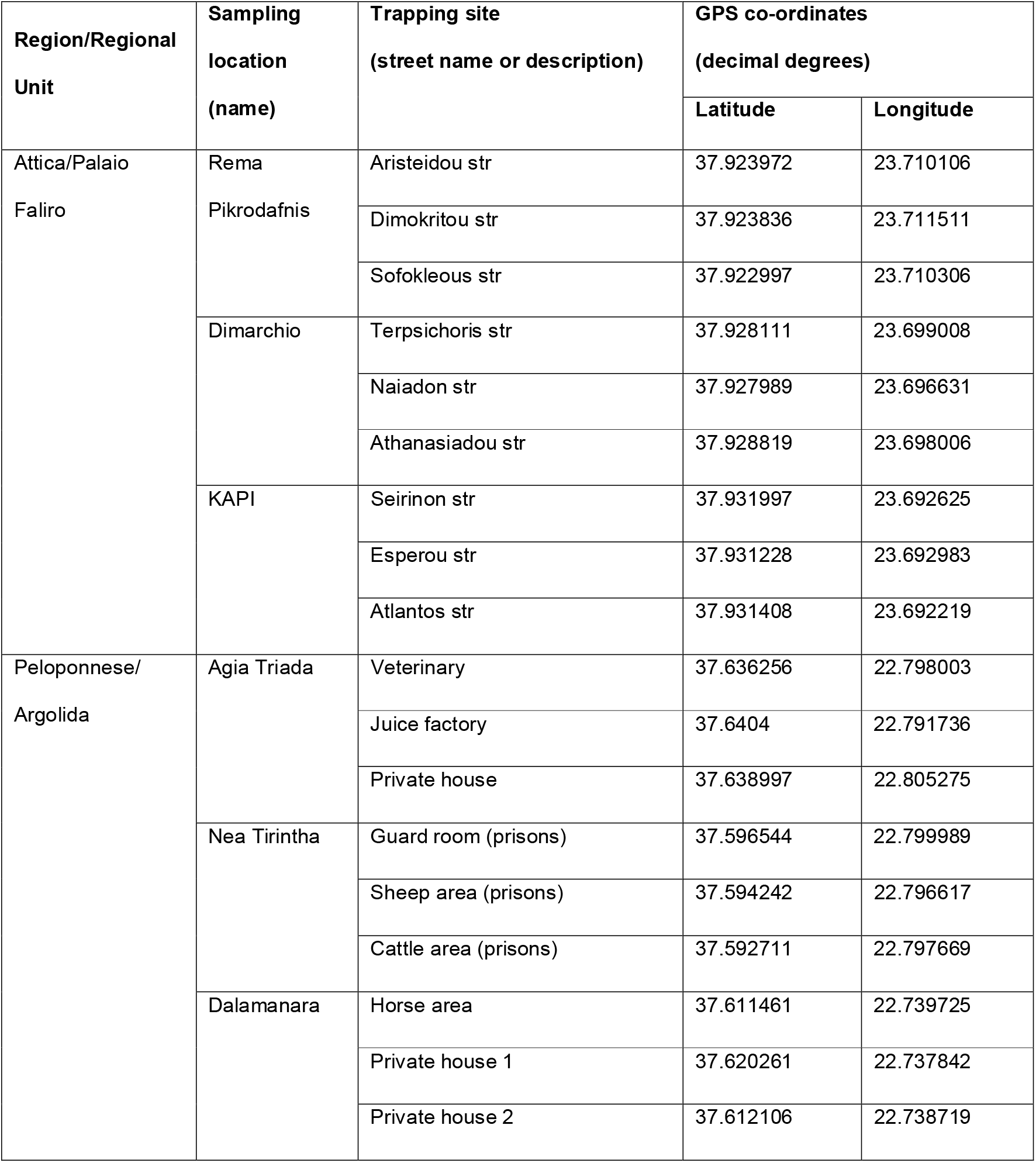
Geographical locations with GPS co-ordinates of mosquito trapping sites within the Attica and Peloponnese regions of Greece.

### DNA/RNA extraction and cDNA synthesis

DNA was extracted from individual male mosquitoes using QIAGEN DNeasy Blood and Tissue Kits according to manufacturer’s instructions. DNA extracts were eluted in a final volume of 100 μL and stored at −20°C. RNA was extracted from individual female mosquitoes using Roche High Pure RNA Isolation Kits and QIAGEN RNeasy 96 kits according to manufacturer’s instructions. RNA extracts were eluted in a final volume of 45 μL and stored at −80°C. RNA was reverse transcribed into complementary DNA (cDNA) using an Applied Biosystems High Capacity cDNA Reverse Transcription kit. A final volume of 20 μL contained 10 μL RNA, 2 μL 10X RT buffer, 0.8 μL 25X dNTPs (100 mM), 2 μL 10X random primers, 1μL reverse transcriptase and 4.2 μL nuclease-free water. Reverse transcription was undertaken in a Bio-Rad T100 Thermal Cycler as follows: 25°C for 10min, 37°C for 120min and 85°C for 5min, with the cDNA stored at −20°C.

### Molecular identification of species

Specimens morphologically identified as within the *Cx. pipiens* complex were identified to species level using a combination of multiplex species-specific PCR assays [34,35]. Additional confirmation of species was undertaken using sequencing of conserved cytochrome c oxidase 1 (*CO1*) gene fragments [36–38]. PCR products were separated and visualized using 2% E-gel EX agarose gels (Invitrogen) with SYBR safe and an Invitrogen E-gel iBase Real-Time Transilluminator. PCR products were submitted to Source BioScience (Source BioScience Plc, Nottingham, UK) for PCR reaction clean-up, followed by Sanger sequencing to generate both forward and reverse reads. Sequencing analysis was carried out in MEGA7 [39] as follows. Both chromatograms (forward and reverse traces) from each sample was manually checked, analyzed, and edited as required, followed by alignment by ClustalW and checking to produce consensus sequences. Consensus sequences were used to perform nucleotide BLAST (NCBI) database queries and sequences were compared to those available from GenBank (NCBI). Representative full consensus sequences for *CO1* gene fragments were submitted to GenBank and assigned accession numbers MN005042-MN005056.

### WNV screening

WNV detection was undertaken using a WNV-specific real-time PCR assay [40]. Reactions were prepared using 5 μL of Qiagen QuantiTect SYBR^®^ Green Master mix, a final concentration of 1 μM of each primer, 1 μL of PCR grade water and 2 μL template cDNA, to a final reaction volume of 10 μL. Prepared reactions were run on a Roche LightCycler^®^ 96 System and PCR cycling conditions were as follows: 95°C for 10 min followed by 45 cycles of 95°C for 10 sec, 60°C for 10 sec, 72°C for 20 sec. PCR products were also separated and visualised using 2% E-Gel EX agarose gels (Invitrogen) with SYBR safe and an Invitrogen E-Gel iBase Real-Time Transilluminator to confirm successful amplification of the 144 base pair target fragment.

### WNV case mapping

Maps were constructed in ArcMap 10.5 (Esri, ArcGIS) using Global Administrative layers for Greece (level 3), downloaded from www.gadm.org (Version 3.6) and anonymized ECDC WNV case report data from “Transmission of West Nile virus, June to December 2018 – Table of cases, 2018 transmission season” downloaded from www.ecdc.europa.eu. The EU NUTS (Nomenclature of territorial units for statistics) level 3 regions as listed in the ECDC data sheet were matched to the Global Administrative layers level 3 (municipalities) during map construction, with each of the GADM level 3 municipalities matched to the corresponding NUTS level 3 region and assigned the same reported case data. The data from the ECDC surveillance Atlas was collected for each week of the transmission season, for human and equine cases, and then combined for each region, to generate maps of monthly reports.

### Statistical analysis

Non-parametric Mann Whitney U tests were performed in Microsoft Excel (version 16.21.1) to compare the number of *Cx. pipiens* complex mosquitoes for each trap type in a given sampling location.

## Results

### Mosquito species abundance and diversity

A total of 1062 mosquitoes comprising 840 unfed females, 28 blood fed females, 9 gravid females and 185 males were captured (**Table 2**). Species belonging to the *Cx. pipiens* complex were the most abundant, comprising 62.5% (n= 664) of the total collection across both RUs. Additional species collected included *Cs. longiareolata* (16.1%, n= 171), *Ae. caspius* (11.0%, n=117), *Ae. albopictus* (7.4%, n=79) and species belonging to the *Anopheles (An.) maculipennis* complex (1.8%, n=19). The remaining 1.1% (n=12) of mosquitoes were not possible to morphologically identify using keys due to damage during trapping. Individuals of the *Cx. pipiens* complex *and Cs. longiareolata* were collected from all sites within both regions. In the Attica region, *Ae. albopictus* was collected in all three sites and single individuals were also collected in Agia Triada and Dalamanara within the RU of Argolida. In contrast, *Ae. caspius* and *An. maculipennis* complex individuals were collected in all three sites within the RU of Argolida but not from sites within the RU of Palaio Faliro.

**Table 2.**
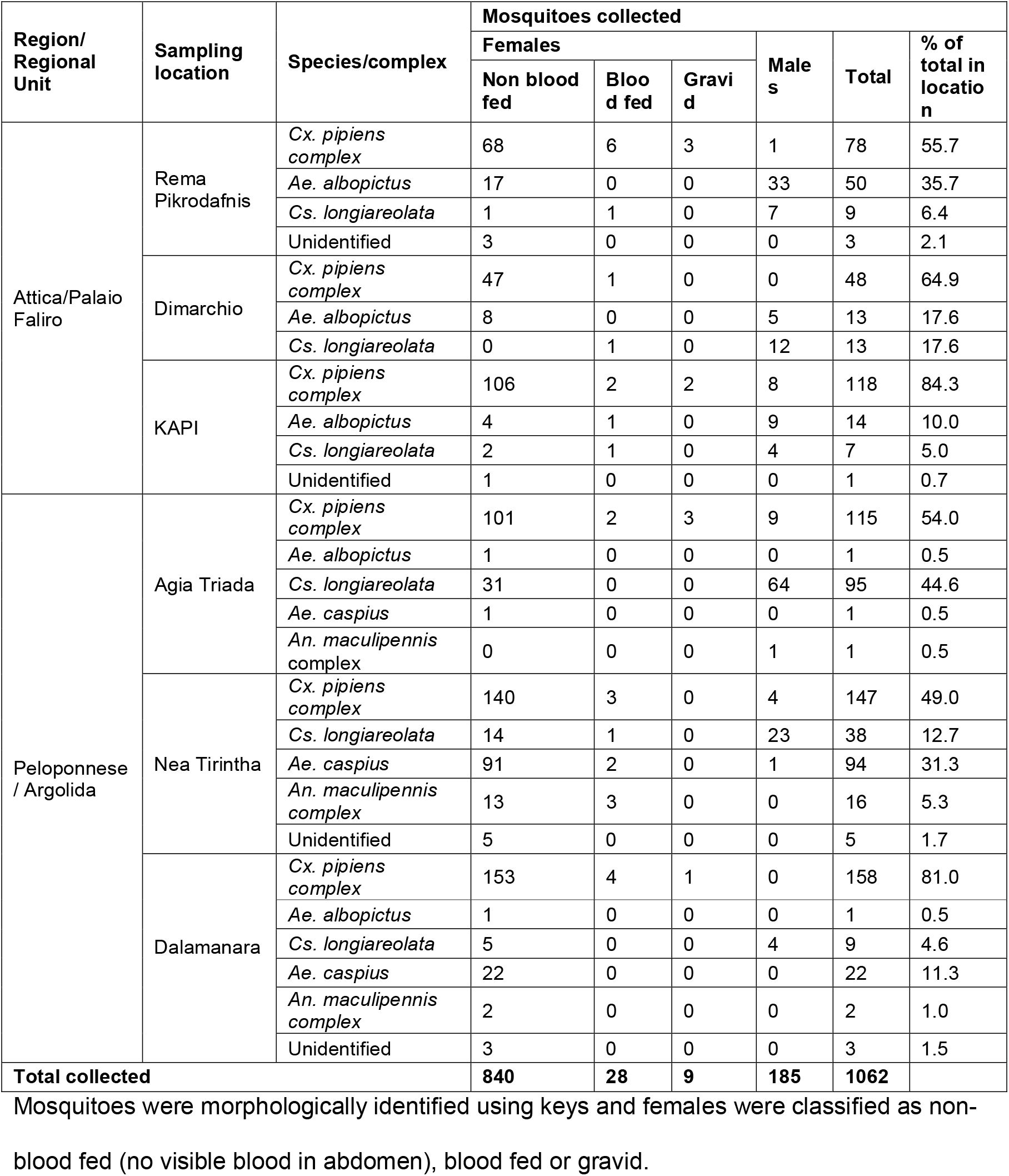
Mosquitoes collected from different locations in the Attica and Peloponnese regions of Greece using a variety of mosquito trap types.

### Species trap comparison

In both RUs, BG traps collected both more overall mosquitoes of all species, and a greater number of specimens from the *Cx. pipiens* complex than CDC traps and EVS traps (**Table 3**). As the data was not normally distributed, non-parametric Mann-Whitney tests were used to determine any significant differences in the number of *Cx. pipiens* complex mosquitoes collected using different trap types (**Table 3**). In the RU of Palaio Faliro, BG traps collected more *Cx. pipiens* complex mosquitoes (n=101) than CDC (n=46) and EVS (n=41) traps although the comparison between BG and CDC traps was not statistically significant (Mann-Whitney U=258.0, p=0.07). In the RU of Argolida BG traps collected significantly more *Cx. pipiens* complex (n=214) than CDC (n=69) and EVS (n=50) traps (Mann-Whitney U=40, p=0.02; U=32, p=0.01 respectively).

**Table 3.**
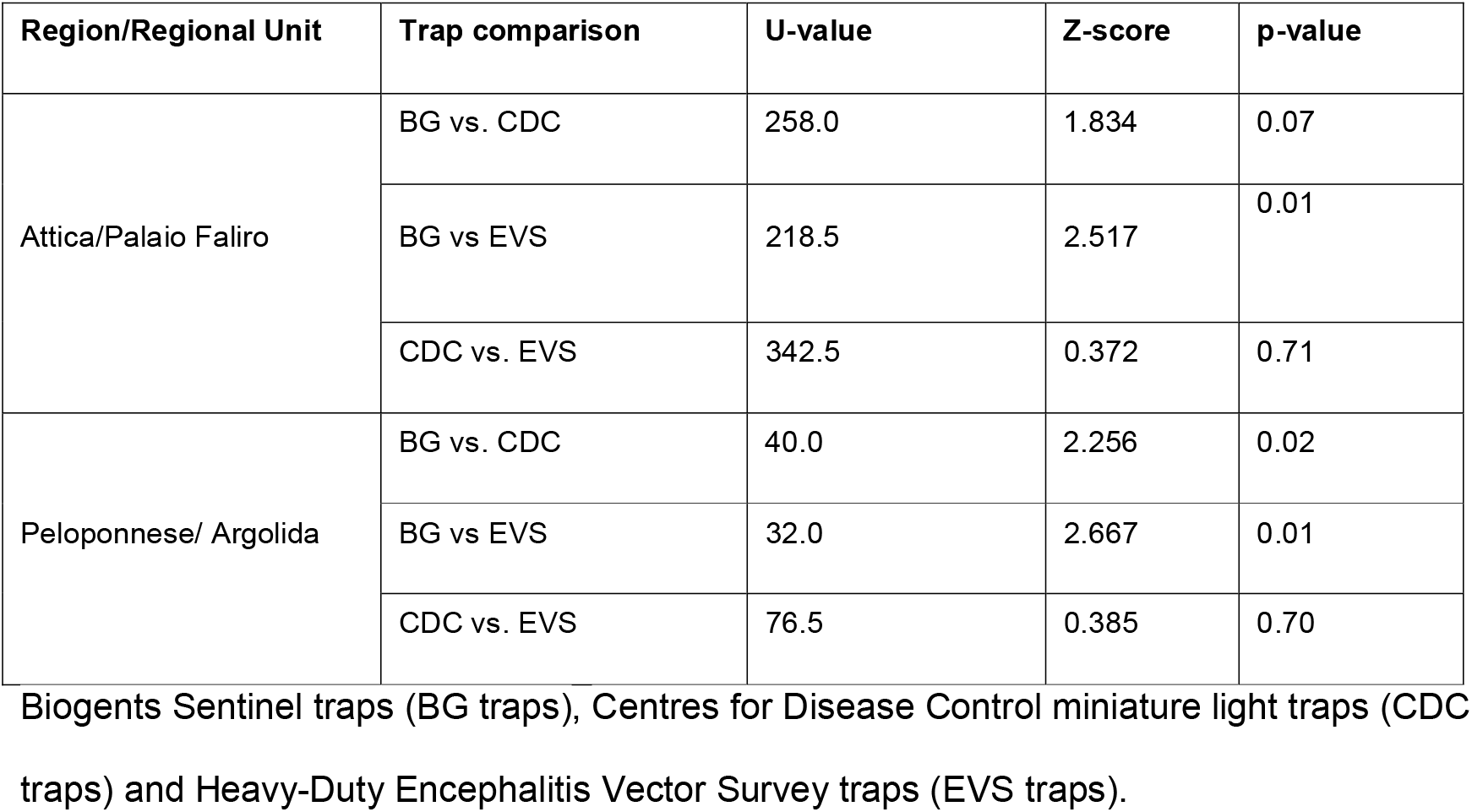
Mann-Whitney statistical analysis comparing the number of *Cx. pipiens* complex mosquitoes collected using three traps.

### Molecular identification of species

Sanger sequencing of *CO1* gene fragments was undertaken to confirm morphological identification of species and to also determine the species of unidentified specimens that had been damaged during trapping. Representative *CO1* gene fragment sequences from individuals of the *Cx. pipiens* complex from all six collection sites across both RUs were obtained using a PCR assay designed for species identification for European *Cx. pipiens* complex species (38) (**Table 4**).

**Table 4.**
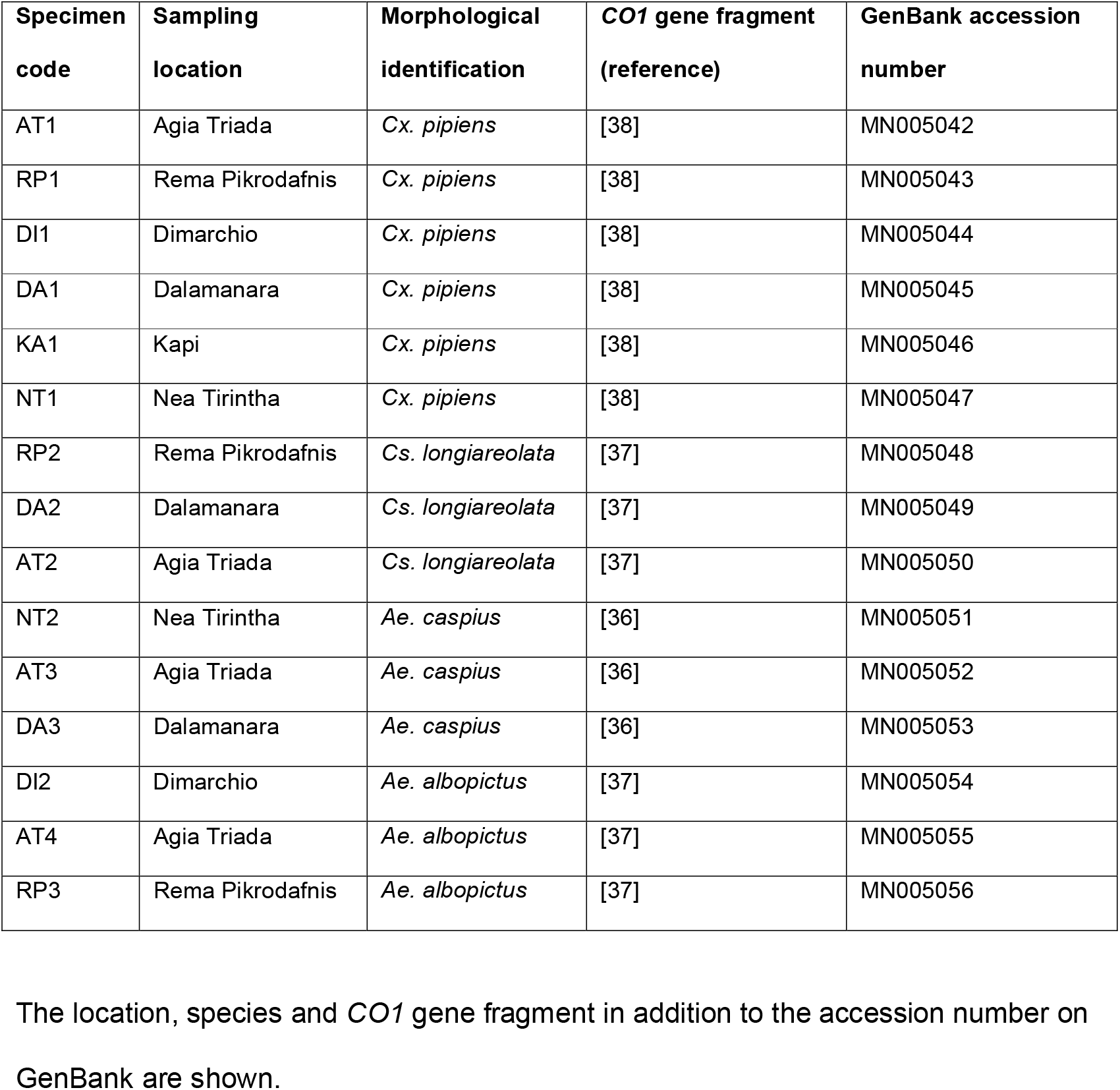
*CO1* GenBank accession numbers for representatives of species confirmed by molecular identification.

Sequencing an additional *CO1* fragment [37] successfully confirmed the identification of *Cs. longiareolata* (n=3) and *Ae. albopictus* (n=3). Sequencing of a third *CO1* fragment [36] was required to successfully confirm *Ae. caspius* (n=3). However, Sanger sequencing of both *CO1* and internal transcribed spacer – 2 (*ITS2*) fragments [41] did not produce sequences of sufficient quality to successfully speciate individuals morphologically identified as within the *An. maculipennis* complex. Multiplex species-specific assays [42,43] revealed the presence of both biotypes of *Cx. pipiens (pipiens* type and *molestus* type) in addition to hybrids (**Fig. 2**). In the RU of Palaio Faliro overall 54.5% (n=79) were confirmed as the *pipiens* type, 20.0% (n=29) as the *molestus* type and 25.5% (n=37) as hybrids. In the RU of Argolida, 68.1% (n=98) were *pipiens* type, 8.3% (n=12) *molestus* type and 23.6% (n=34) hybrids.

**Fig. 2.**
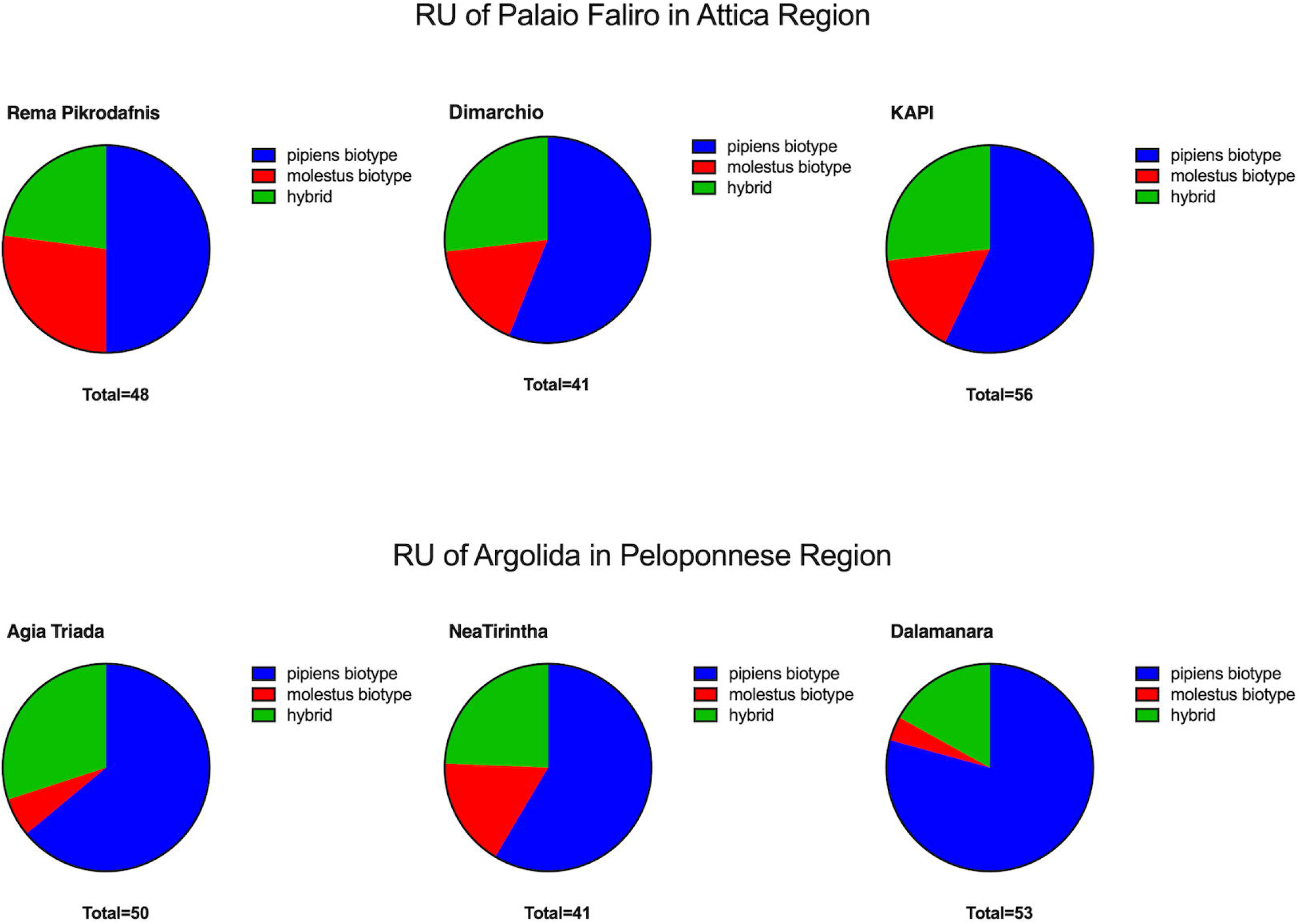
Prevalence rates of *Cx. pipiens* biotypes. Mosquitoes analysed using multiplex species-specific PCR assays were from three sampling locations in Regional Unit of Palaio Faliro in Attica Region (A) and Regional Unit of Argolida in Peloponnese region (B) of Greece during May-June 2018.

### WNV infection rates in field mosquitoes

A total of 630 individual mosquitoes (229 from RU of Palaio Faliro and 401 from RU of Argolida) were screened for the presence of WNV RNA and four *Cx. pipiens* complex individuals were WNV positive. qRT PCR results were confirmed by separation and visualisition of PCR products using gel electrophoresis. These positive individuals were unfed females which were molecularly identified as three *pipiens* type and one hybrid, all collected from the RU of Argolida at the end of May. This is interesting when compared to the spatial and temporal records of human and equine cases during 2018 (**Fig. 3**) as only one human case was recorded from the Peloponnese region all year, and not until August. This is in contrast to the RU of Palaio Faliro where, across the whole Attica region a total of 159 human cases were recorded in 2018, with the first reported cases occurring in June, but no WNV was detected in the mosquitoes collected from this region.

**Fig. 3.**
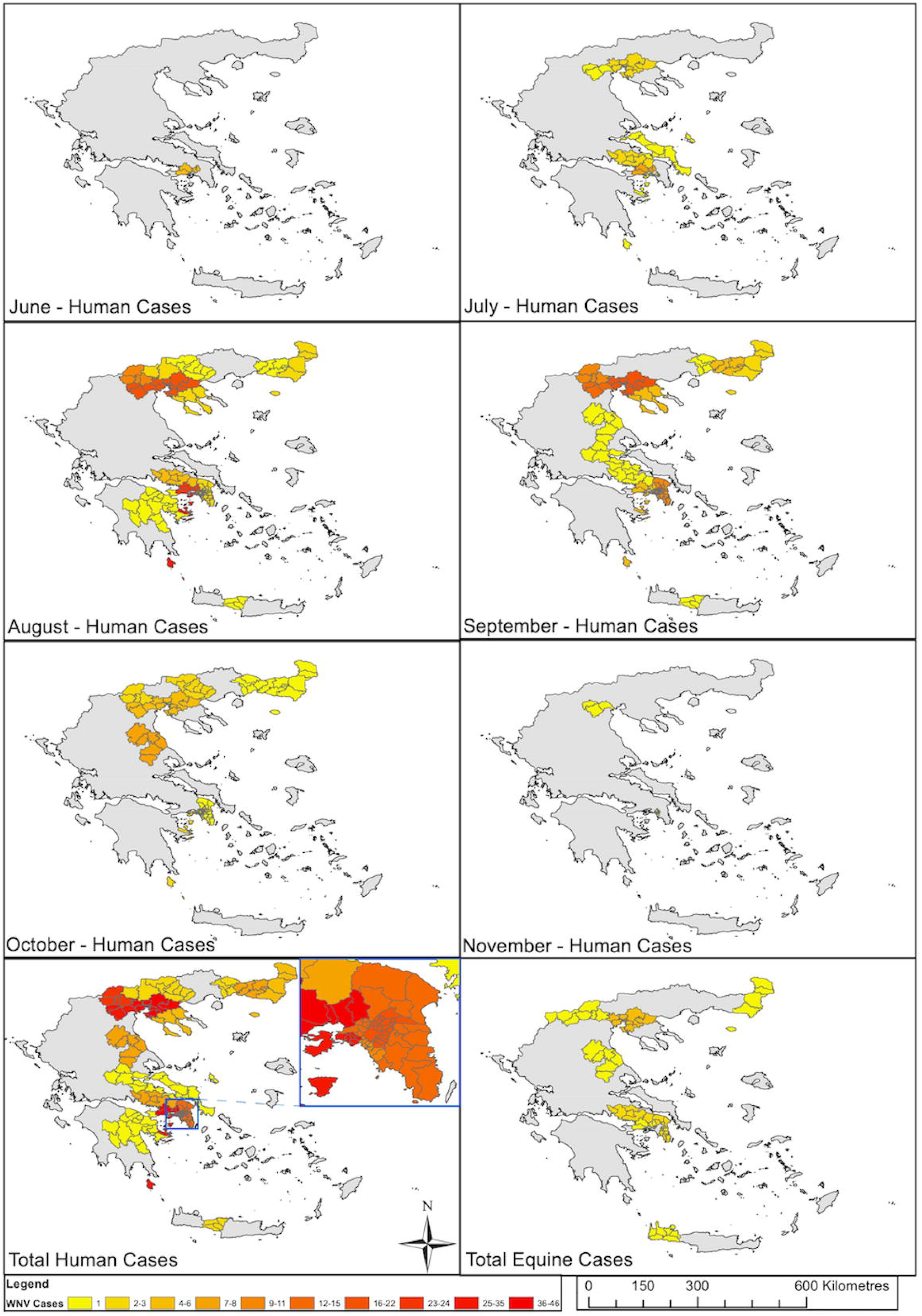
Reported human and equine cases of WNV in the 2018 transmission season. Maps were constructed in ArcMap 10.5 (Esri, ArcGIS) using Global Administrative layers for Greece (level 3), downloaded from www.gadm.org (Version 3.6) and ECDC WNV case report data from “Transmission of West Nile virus, June to December 2018 – Table of cases, 2018 transmission season” downloaded from www.ecdc.europa.eu. The data from the ECDC surveillance Atlas was collected for each week of the transmission season, for human and equine cases, and then combined for each region, to generate maps of monthly reports.

## Discussion

Our mosquito trapping experiments using different adult traps show that in both regions BG traps collected both a larger number of mosquitoes of all species, and a greater number of individuals from the *Cx. pipiens* complex (although this was not statistically significant in the RU of Palaio Faliro). Previous trap comparison studies undertaken in Europe report contrasting results, ranging from BG traps in Germany collecting more *Cx. pipiens* complex mosquitoes than CDC and EVS traps [44], to a study in Spain showing no statistically significant differences between BG and CDC traps in collecting specimens from this complex [45]. Although we measured temperature and humidity during our collection periods (Additional file 1), there are a variety of additional factors that can influence the collections obtained from adult mosquito traps including wind and the use of different attractants. Our results highlight that using a variety of trapping types can increase the species diversity of collections, however, targeting resources to just use BG traps may enable a greater number of target vector species – such as individuals of the *Cx. pipiens* complex – to be collected.

Although different mosquito species (across multiple genera) have been demonstrated to be competent vectors of WNV [46], the major vectors for WNV belong to the *Cx. pipiens* complex. In this study, we collected individuals of the *Cx. pipiens* complex in addition to other species including *Ae. albopictus, Cs. longiareolata* and *Ae. caspius* shown previously to be present in Greece [47,48]. The presence of the *pipiens* type, *molestus* type and hybrids in both the Attica and Peloponnese regions is consistent with previous studies in Greece [22,49,50]. We found variation in the prevalence of the different types with the *pipiens* type comprising 54.5% (n=79) in the RU of Palaio Faliro, 20.0% (n=20) *molestus* type and 25.5% (n=37) of hybrids. These results differ from another study that had found a more homogeneous *molestus* type population [49]. In RU of Argolida the biotypes of the *Cx. pipiens* complex were 68.1% (n=98) of *pipiens* type, 8.3% (n=12) of *molestus* type and 23.6% (n=34) of *molestus* and *pipiens* hybrids. The high percentage of hybrids in the RU is similar to a previous study conducted in the area after the 2017 outbreak that reported 37% hybrids, 41% *pipiens* and 22% *molestus* types [50].

The two biotypes are morphologically indistinguishable but have genetic, biological and behavioural differences. The *pipiens* type is anautogenous, so females need to consume a blood meal to lay eggs [22]. Furthermore, the *pipiens* type requires a large space to swarm for mating, are found above ground undergoing diapause and are primarily ornithophilic (preferring to feed on birds). In contrast, the *molestus* type is autogenous and can lay eggs without a blood meal. Mating can happen in confined spaces, while they live underground, do not undergo diapause, and are more anthropophilic, preferentially feeding on humans. Hybrid types are important in the epidemiology of WNV. In the USA, the high number of WNV cases in humans was correlated to the high number of hybrids [51]. Europe is considered to have more “pure” types but hybridization can result in a catholic feeding behaviour (feeding both on birds and mammals) increasing the risk of mixed populations acting as bridge-vectors of WNV between birds and humans/equines [49]. The feeding patterns of the different mosquito species, and the different types within the species complex, are important in order to identify the contribution of each vector to both the enzootic maintenance of WNV in avian hosts, and the spill-over transmission to humans and horses [52]. In northern Greece, the predominance of *pipiens* type could be facilitating the maintenance of the enzootic cycle of the virus between mosquitos and birds in the area [49]. The presence of the *molestus* type and the existence of hybrids can promote an opportunistic biting behaviour that could contribute to the spill-over of infection to humans and equines.

In our study, we also collected several other species that have been implicated or shown to be potential WNV vectors. Experimental transmission has been shown for both *Cs. longiareolata and Ae. albopictus* [1]. Species belonging to the *An. maculipennis* group are considered potential vectors of WNV [2]. Laboratory experiments have indicated that *Ae. caspius* may be incapable of transmitting WNV [27,53], however, in some countries the high densities, and detection of WNV in wild-caught specimens, have suggested this species may have a potential role in transmission, particularly during an outbreak when the level of virus circulation is high [54]. The presence of *Ae. albopictus*, an invasive species that has expanded its range across Europe since the late 1970s, would suggest the potential for transmission of additional arboviruses. *Ae. albopictus* has the ability to adapt to colder temperatures and stay dormant during the winter and has previously been shown to be responsible for chikungunya virus outbreaks in Italy in 2007 [55]. *Ae. albopictus* has also been the principle vector responsible for dengue virus outbreaks in Hawaii in 2001-2002 and Mauritius in 2009 [56,57] and is a potential vector of Zika virus [58,59]. Furthermore, it can be a competent vector of WNV when experimentally tested in laboratory conditions [60] and in North America natural infections have been found. The opportunistic biting behaviour of *Ae. albopictus* may increase this species role as a vector of WNV. In Greece, since its first reported presence in 2003 in the western part of the country, *Ae. albopictus* has now spread to almost every district [48].

Detection of WNV virus RNA in four unfed *Cx. pipiens* complex specimens would indicate circulation of WNV in the RU of Argolida during our collection period in May. This would be supported by one human laboratory confirmed case reported in Peloponnese in 2018, however, it is interesting to note the reported human case didn’t occur until August, suggesting WNV may have been circulating in the area for months before resulting in a case of human clinical disease. The confirmation that three of the positives were *pipiens* type, supports the possibility of virus circulating in an enzootic cycle, between birds and mosquitoes. However, the presence of WNV in one of the hybrids also demonstrates the potential for spill-over transmission to humans and equines in the area at this early time in the season. In comparison, no WNV was detected in mosquitoes collected from the RU of Palaio Faliro, but this area subsequently recorded a far greater number of human and equine cases during 2018, highlighting the likely variations in spatial and temporal transmission dynamics between these two very different localities, and the variable factors that can influence risk of infection and disease during the transmission season.

## Conclusions

Sampling during the onset of the 2018 WNV transmission season in the RUs of Attica and Peloponnese Regions was particularly important in a year in which more than 300 human cases were recorded in Greece. These results, combined with previous entomological surveys conducted in Greece, show the high occurrence of hybrids between the *pipiens* and *molestus* types of *Cx. pipiens*. Previous studies have demonstrated the importance of hybrids as bridge vectors of WNV. Their role in spill-over transmission to humans, and the presence of hybrids (and WNV infections) in RUs of Attica and Peloponnese regions of Greece suggest these areas are vulnerable to outbreaks. Furthermore, 2018 was the first year in Greece in which WNV human cases were recorded so early in the transmission period with six human cases confirmed by late June. Future entomological surveillance studies should incorporate molecular xenomonitoring to determine this potential expansion of the transmission season to provide early warning systems for potential WNV outbreaks. Notification of human WNV cases in Europe through the The European Surveillance System (TESSy) [61] of the ECDC allows a weekly map of human cases [31]. In addition, reporting of WNV encephalomyelitis in horses to the European Commission is carried out via the Animal Disease Notification System (ADNS). As reported cases of WNV infection in humans have been from southern and central European countries and a majority of human infections are asymptomatic, it is particularly important to undertake entomological and avian surveillance to determine if WNV circulation is occurring in particular area. In particular, entomological surveys to determine the distribution of mosquito vectors such as *Cx. pipiens* through the Pan-European VectorNet [62] will play a crucial role in an integrated approach to WNV surveillance and control efforts to minimise the impact of outbreaks on veterinary and public health.

## List of abbreviations

WNV: West Nile virus
*Cx*: *Culex*
RUs: Regional Units
BG traps: Biogent sentinel traps
CDC traps: Centres for Disease Control miniature light traps
EVS traps: Heavy Duty Encephalitis Vector Survey traps
*Ae*: *Aedes*
*Cs*: *Culiseta*
*An*: *Anopheles*
ECDC: European Center for Disease Control
*CO1*: cytochrome c oxidase 1

## Declarations

### Ethics approval and consent to participate

The study protocol was reviewed and approved by the the institutional review boards of the London School of Hygiene and Tropical Medicine (#15234).

### Consent for publication

Not applicable.

### Availability of data and materials

All representative mosquito species sequences are available from Genbank: accession numbers MN005042-MN005056. The datasets generated on Collection, extraction and PCR results are available at Open Science Framework: DOI 10.17605/OSF.IO/D76QF.

### Authors’ contributions

MB contributed to conceptualization, data curation, formal analysis, investigation, methodology and writing of the original draft. CLJ contributed to conceptualization, data curation, formal analysis, investigation, methodology, supervision, writing of the original draft and review and editing of the manuscript. IL contributed to investigation and methodology. AM contributed to conceptualization, investigation, methodology, project administration, supervision and review and editing of the manuscript. TW contributed to conceptualization, data curation, formal analysis, investigation, methodology, funding acquisition, supervision, project administration, supervision, writing of the original draft and review and editing of the manuscript. All authors read and approved the final manuscript.

## Acknowledgements

The authors would like to thank Mrs A. Fikiri, Mrs A. Georgopoulou (Municipality of Palaio Faliro), Mr V. Karras, Mr G. Balatsos (Benaki Phytopathological Institute) and Mr P. Kalkounos (Agricultural Prison of Tirintha) for their active help to place the trap networking. We also want to thank Prof. D. Petric and Dr D. Papachristos for their comments and suggestions for the methodology on vectors surveillance.

## Additonal file 1

Temperature (°C) and relative humidity (%) during the collection periods in Regional Unit of Palaio Faliro in Attica Region (A) and Regional Unit of Argolida in Peloponnese region (B) of Greece.

