## Supplementary figures and images for "The abundance and diversity of West Nile virus mosquito vectors in two Regional Units of Greece during the onset of the 2018 transmission season"

### additional file 1

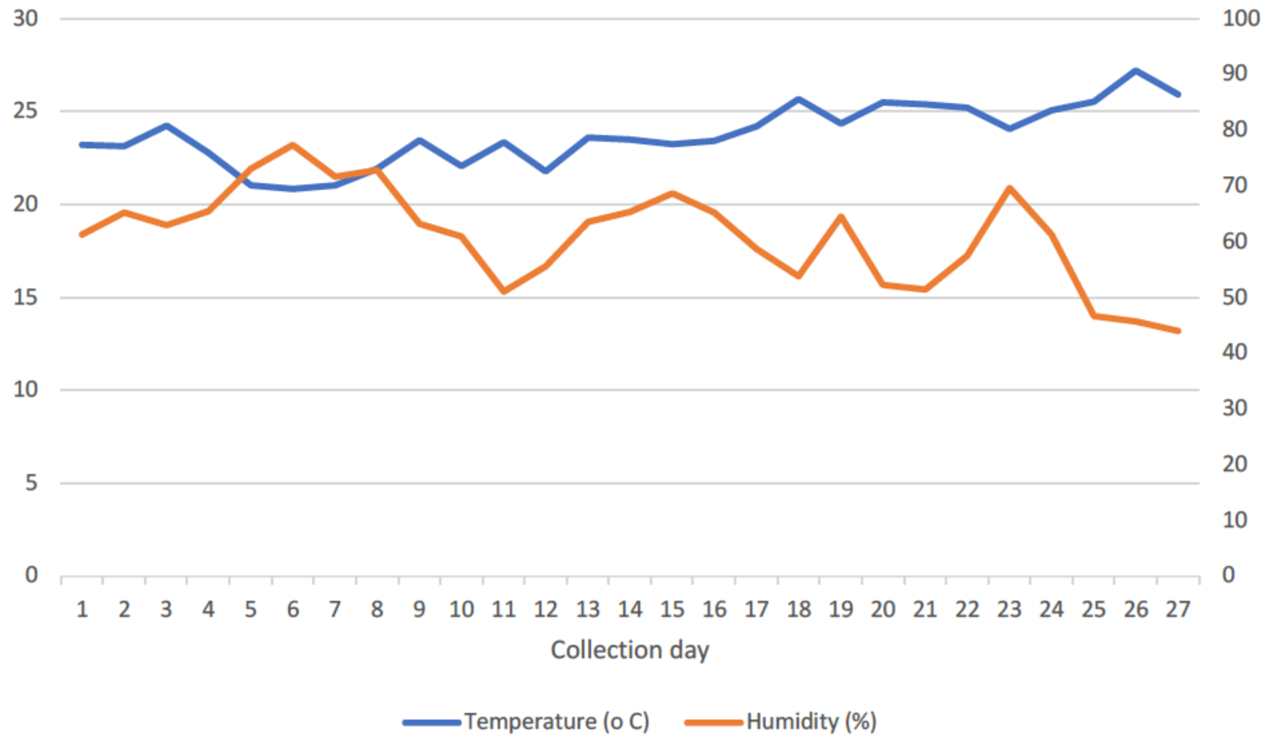

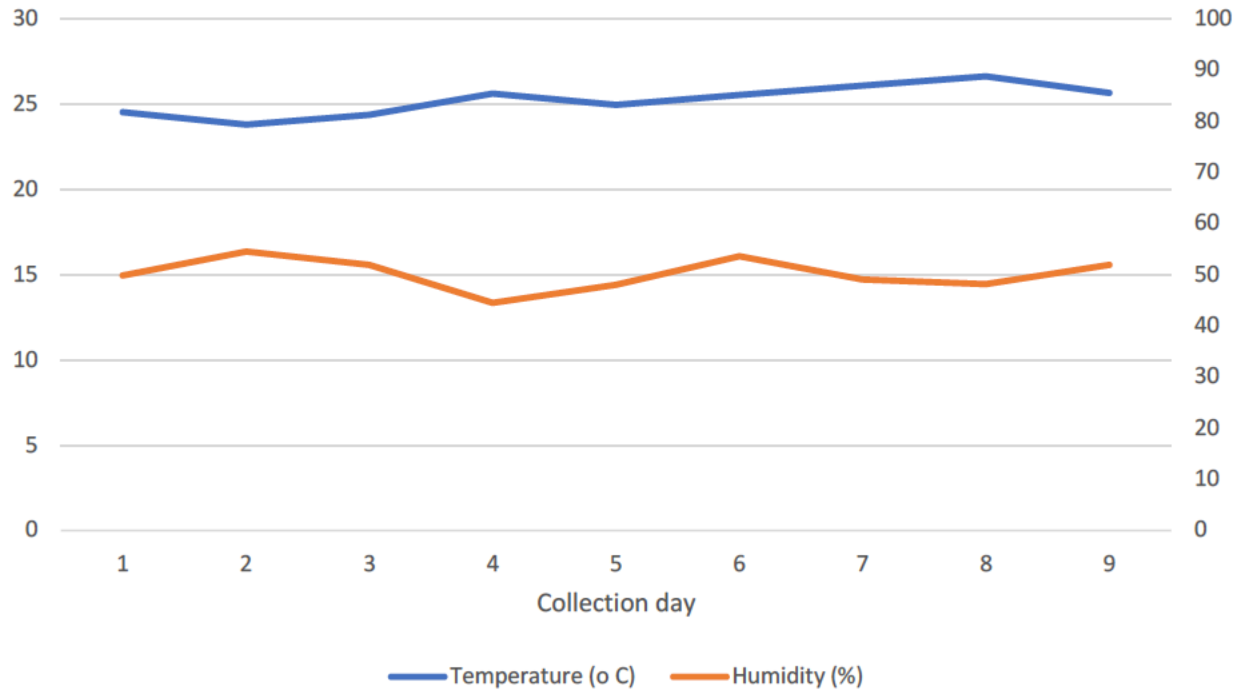
